# A honey authenticity test by interphase emulsion reveals biosurfactant activity and biotechnology in the stingless bee nest of *Scaptotrigona* sp. ‘Catiana’ from Ecuador

**DOI:** 10.1101/2022.05.11.491040

**Authors:** Patricia Vit

**Affiliations:** Apitherapy and Bioactivity, Food Science Department, Faculty of Pharmacy and Bioanalysis, Universidad de Los Andes, Mérida, Venezuela

**Keywords:** honey authenticity test by interphase emulsion HATIE, honey biosurfactant test HBT, pollen residue, Roquefort cheese smell, *Scaptotrigona*, sensory evaluation, stingless bee honey

## Abstract

Bees are valuable pollinators of fruit trees and grasses. Honey is a medicinal food of plant and animal origin, with social impact for the wellbeing of bee keepers. The Neotropical apifauna has about 500 species of stingless bees (Meliponini). Commercial beekeeping with *Apis mellifera* underestimates the cultural legacy of meliponiculture, and both are affected by the presence of fake honeys in the market. Three alternative techniques (interphase emulsion, sensory analysis, and pollen residue) to nuclear magnetic resonance (NMR) were investigated to detect false honeys. One technique was based on an interphase emulsion test, which can be performed by beekeepers, farmers, housekeepers, professionals and consumers of honey in general. Another technique was sensory analysis by a tasting panel, and the third consisted on a palynological preparation with a simplified observation. Five genuine honeys produced by *Apis mellifera, Geotrigona leucogastra, Melipona mimetica, Scaptotrigona* sp., *Tetragonisca angustula* and a fake honey from Ecuador were analyzed. The honey authenticity test by interphase emulsion was fast and effective to detect fake honey (two phases), and genuine honeys (one and three phases). A further screening of was done in 51 Asian, Australia, European and Latinamerican honeys. Additionally the HATIE generated a new application as a test to detect unique biosurfactants in honey (HBT) of *Scaptotrigona* sp. ‘Catiana’ (one phase) with potential microbial origin, and its entomological origin in this set of honeys. ‘Catiana’ nests smell like Roquefort cheese, indicating a fungus association with this rural stingless bee highlighted by its distribution, productivity and the peculiarities described in this research, 80 years after its description as a new genus *Scaptotrigona* Moure, 1942. Paradoxically, this communication without microbiological analysis, infers the fungal presence in the nest of *Scaptotrigona* sp. by sensory observations different from the classic sensory evaluation of honey.

## 1. Introduction

Neotropical stingless bees comprise 500 of the 600 identified species in the world. They thrive from 27.03°N in North America, Álamos, Sonora, Mexico, up to 34.90°S in South America, Montevideo, Uruguay (Roubik and Vergara, 2021). *Scaptotrigona* Moure 1942 was the first new genus described by Father Jesus Santiago Moure (1912-2010) 80 years ago, the most prolific meliponine taxonomist to date. *Scaptotrigona* is the most abundant stingless bee in Ecuador producing most pot-honey in the country, they are kept by farmers who are familiar with them in the field. Basic knowledge on stingless bee management is transmitted within families. A market study determined that there was a supply of about 500 L of honey and a demand of 2,000 L, for which the feasibility of breeding *Scaptotrigona* in the high zone of El Oro province, Ecuador was proposed (Aguirre Zambrano, 2015). The ethnic name of this bee is ‘Catiana’ in El Oro province, and ‘Catana’ in the Loja province.

Two Kichwa lady sensory assessors from the Rio Chico community, near Puyo, capital of Pastaza province, Amazonas, were extraordinary. They tasted three pot-honeys (Meliponini), two *Apis mellifera* honeys, and a fake honey, which was disgusting (Vit et al., 2017). In contrast with an urban panel of 58 assessors which selected a fake honey as their favorite during the I Congress of Apiculture and Meliponiculture in Machala, Ecuador (Vit et al., 2015a). With these two examples, bonds with the territory became links for ancestral knowledge and sensory perception.

The oldest records of meliponiculture are by the Mayan culture with the stingless bee *Melipona beecheii* called ‘Xunan Kab’ (Weaver y Weaver, 1981), still productive and managed to preserve the Mayan ancestral traditions (Sic México, 2012) for the cosmovision in the Yucatan Peninsula, Mexico (Sotelo Santos & Alvarez Asomoza, 2018). In Venezuela, the Huottuja (Piaroa) from the Amazonas state keep stingless bees of diverse species (Vit et al., 2011) of the *Melipona, Scaptotrigona* and *Tetragona* genera. Ecuador has an apifauna of about 200 stingless bees, which have not reached the Galapagos. Coloma collected 78 species during his master’s thesis (1986). Dr. Silvia R.M. Pedro from the Universidade de São Paulo in Ribeirão Preto, Brazil, identified 54 species of stingless bees collected during a year (Vit et al., 2018), and 100 species of stingless bees come from a tiny 64 km^2^ plot of Yasuní (Roubik, 2018). Brazil has some 250 species of meliponines in a much larger territory (Melo, 2020).

Sustainable bee keeping and meliponiculture allow maintaining the environmental biodiversity of bees and plants, along with medicinal and social benefits for producers, aimed at reducing poverty (FAO, 2021). Its integration into rural development programs in Ecuador to improve well-being (Álvarez Montalvo, 2021) has versatile strategic potential in key aspects of pollination, beehive products and apitherapy. In an organized community with bee keeping and meliponiculture professionals, there is no room for fake honey. With so many floral and entomological resources, honey biodiversity in Latin America is a treasure protected by ancestral knowledge but forgotten by the industry and agricultural development plans, the optimal terrain for the marketing of fake honey. Honey from bees is a product that occupies a place of honor in cupboards, due to its medicinal use (Vit et al., 2015b). Together with pollen, propolis and royal jelly, they are eye-catching products at any agricultural fair where they can be promoted. In addition to its economic impact, Paredes and Saravia (2021) highlighted the function of social and cultural meetings in the farmer fair space to communicate specialized agricultural management.

Recently, the composition of honey from meliponines was reviewed in a context where the global honey market is dominated by *Apis mellifera* (Vit, 2017). The physical-chemical composition of honey from *Apis mellifera* marketed in Quito was analyzed (Velásquez and Goetschel, 2019), and honey from Meliponini stingless bees from Ecuador (Villacrés-Granda et al., 2021), Costa Rica (Umaña et al., 2021), Australia and Malaysia (Zawawi et al., 2022). Nuclear magnetic resonance is a rapid method to confirm the authenticity of honey (Spiteri, 2015; Yong et al., 2022) and its entomological origin (Vit et al., submitted). Since 2014, the Honey Authenticity Test by Interphase Emulsion (HATIE) produced a novel result in honey from *Scaptotrigona* sp., but its possible explanation allowed expanding the application of the authenticity test to a metabolic field of honey biosurfactant activity with suspected microbial origin. Furthermore, the nests of *Scaptotrigona* sp. smell like Roquefort cheese, which is reported for the first time here as a particularity observed in only one of the 54 species of stingless bees collected in Ecuador.

The HATIE was tested on 51 Asian, European and Latin American honeys, to show its universality. When it was published (Vit, 1998), it had already been tested by about a thousand honey consumers (bee keepers, farmers, scientists, housewives, students, teachers, employees and professors, other professions). The Andean bee keeper with hives located at highest altitudes in Venezuela, remembers the “little cloud” as he memorized the emulsion formed by genuine honey more than 30 years ago (Juan Carlos Schwartzenberg, La Casita de la Miel, Escagüey, Mérida, Venezuela, personal communication, 21^st^ April 2022).

In order to value non-traditional techniques to detect fake honey, useful in a scientific and social context for honey consumers, the following was carried out: 1. The honey authenticity test by interphase emulsion (HATIE), 2. Quantitative descriptive sensory analysis, together with sensory acceptance, and 3. Observation of the presence or absence of honey pollen sediment. Thus, decision-making on authenticity was based on interphasic, sensory and microscopic assessments. Genuine honey from *Apis mellifera*, and from stingless bees *Geotrigona leucogastra, Melipona mimetica, Scaptotrigona* sp. and *Tetragonisca angustula*, and a false honey.

## 2. Materials and methods

### 2.1 Honey sampling

Pot-honeys were harvested by suction of pierced cerumen of honey pots. The six honeys analyzed were a genuine commercial *Apis mellifera* honey, four honeys from stingless bees of the most frequent genera in Ecuador *Geotrigona, Melipona, Scaptotrigona* and *Tetragonisca* where these four genera were recommended for the first Pot-Honey Standard Project 2015 http://www.saber.ula.ve/stinglessbeehoney/ during the revision of the Ecuadorian Technical Standard for Honey NTE INEN 1572, in Spanish), and a commercial fake honey recognized for its smell of eucalyptus oil was received from Mr. Julio Vásquez, bee keeper of Pichincha province. During the honey sampling, the nest of *Scaptotrigona* presented the characteristic odor of Roquefort cheese, which was observed in the numerous nests of this bee. A screening with Asian, European, and Latin American honeys, was performed on a collection of 51 frozen honeys from the Apitherapy and Bioactivity Group, Food Science Department.

### 2.2 Identificationn of stingless bees

Stingless bees were collected in isopropyl alcohol, dried and identified by: 1. Prof. Dr. C. Vergara, Department of Chemical and Biological Sciences, Universidad de Las Américas, Puebla, México, as *Geotrigona leucogastra* (Cockerell, 1914) ‘Earth bee’, deposited in the entomological collection UPSE, Peninsula State University of Santa Elena, Ecuador, and 2. Dra. S.R.M. Pedro, Department of Biology, Universidade de São Paulo, Brazil as: 1. *Melipona mimetica* Cockerell 1914 ‘Bermejo’, 2. *Scaptotrigona* sp. ‘Catiana’ and 3. *Tetragonisca angustula* (Latreille 1811) ‘Angelita’, deposited in the Camargo RPSP Collection, Ribeirão Preto, São Paulo, Brazil. Amonth ago the new species *Scaptotrigona ederi* Engel 2022 was recognized (Engel, 2022a described on pp. 6-12), but this species is not the Schwarz *nomina nuda*, it is known that the Schwarz and Engel species are not conspecific with the Ecuadorian ‘Catiana’. The new Ecuadorian species is already characterized and will soon be published (Michael S. Engel, personal communication, 2^nd^ May 2022), which is also deposited in the Snow Entomological Collection, University of Kansas Natural History Museum, Lawrence, Kansas, USA (SEM).

### 2.3 Honey Authenticity Test by Interphase Emulsion (HATIE)

The test to evaluate the authenticity of honey by forming an emulsion as an intermediate phase between the aqueous and the ethereal phases (Vit, 1998), has a patented prototype (Vit, 1995, 1999). A glass vial with a bakelite lid was used, 1.0 g of honey was weighed, which was diluted with 1.0 g of distilled water, then 2.0 mL of ethyl ether were added and it was strongly shaken, left undisturbed for 1 min to allow separation of phases. False honeys form two phases and genuine honeys form three phases, with an emulsion as an intermediate phase, this allows visual detection of false honeys. Except for the genuine honey of *Scaptotrigona*, with a single phase, which is the novel result of the present investigation.

### 2.4 Sensory evaluation of honey

An acceptance test was conducted with 30 assessors (14 women, 16 men, 19-48 years), with a 0-10 cm unstructured line anchored with the expression ‘I like it a little’ 1 cm from the left end, and ‘ I like it a lot’ at 9 cm from the far left (Hein et al., 2008; Lawless and Heymann|, 2010). For this, each honey was served in a glass cup (Gonnet and Vache, 1984) coded with three random digits, each advisor received 6 plastic spoons, a bottle of mineral water, a napkin and a form to mark their acceptance. A quantitative descriptive sensory system was also used, for which a panel of five people (3 women and 2 men, 21-37 years old) tasted the honeys twice, noted their taste, smell and aroma attributes, quantified them and some were graphed. Each sample of honey (5.0 ± 0.1 g) was served in opaque plastic containers with lids, coded with three random digits, served at room temperature on a tray, with a plastic spoon, a napkin, a glass of water and a piece of green apple to consume at the end of each tasting, according to Vit (2008b). The odor-aroma (O-A) description of each honey was made using the 8 families (floral-fruity, vegetable, fermented, wood, nest, honeyed, primitive, industrial chemical), with their corresponding subfamilies and sensory descriptors. The intensity scale 0-3 corresponds to 0 (absent), 1 (mild), 2 (medium), 3 (strong). This intensity scale was also used to quantify persistence and the tastes: acid, bitter, astringent, sweet, pungent, salty, and umami. Likewise, the odor/aroma intensity relationship was presented. A note to indicate that astringent and spicy are trigeminal sensations, since they do not have taste buds for their perception. They are presented as tastes because they are perceptions in the mouth, which facilitates communication with the evaluation panel due to the non-specialized language that uses spicy taste in commercial foods. There may be other sensations in the mouth, rarely parasympathetic salivation, others such as numbness of the tongue. The presence of occasional phytochemicals in honey interacts with our human physiology and causes these sensations.

### 2.5 Pollen sediment of honey

For this analysis, a sediment of the diluted honey was prepared following the melissopalynological method (Louveaux et al., 1978), 10 g of honey were diluted with 20 mL of distilled water in a 100 mL beaker and centrifuged in two plastic conical tubes with lids at 3500 rpm for 10 minutes. The supernatant was discarded, resuspended with 10 mL of water, and centrifuged again. The sediment was spread over an area of 2 × 2 cm^2^ on a microscope slide, and allowed to dry, a drop of glycerine gelatin was added, flattened with a cover slip, and the edges were sealed with transparent enamel. It was observed with a light microscope at X400.

## 3. Results

Genuine pot-honey usually have higher moisture content and free acidity than *Apis mellifera* honeycomb honey. The diastase enzymatic activity is very low in *Melipona* honeys, perhaps for this reason some species better preserve the smell and aroma of the flowers, especially *Melipona favosa* from Venezuela and *Melipona subnitida* from Brazil. Pot-honeys also vary according to the botanical origin but the entomological origin prevails on the nectar variations. It means, among other behavioral variables, the processing in the honey pots inside the nest is different. The microbial associations with stingless bees and their functions for the colony are dawning research challenges. Table 1 presents the results obtained for the six honey types tested here with the three analytical approaches.

**Table 1.**
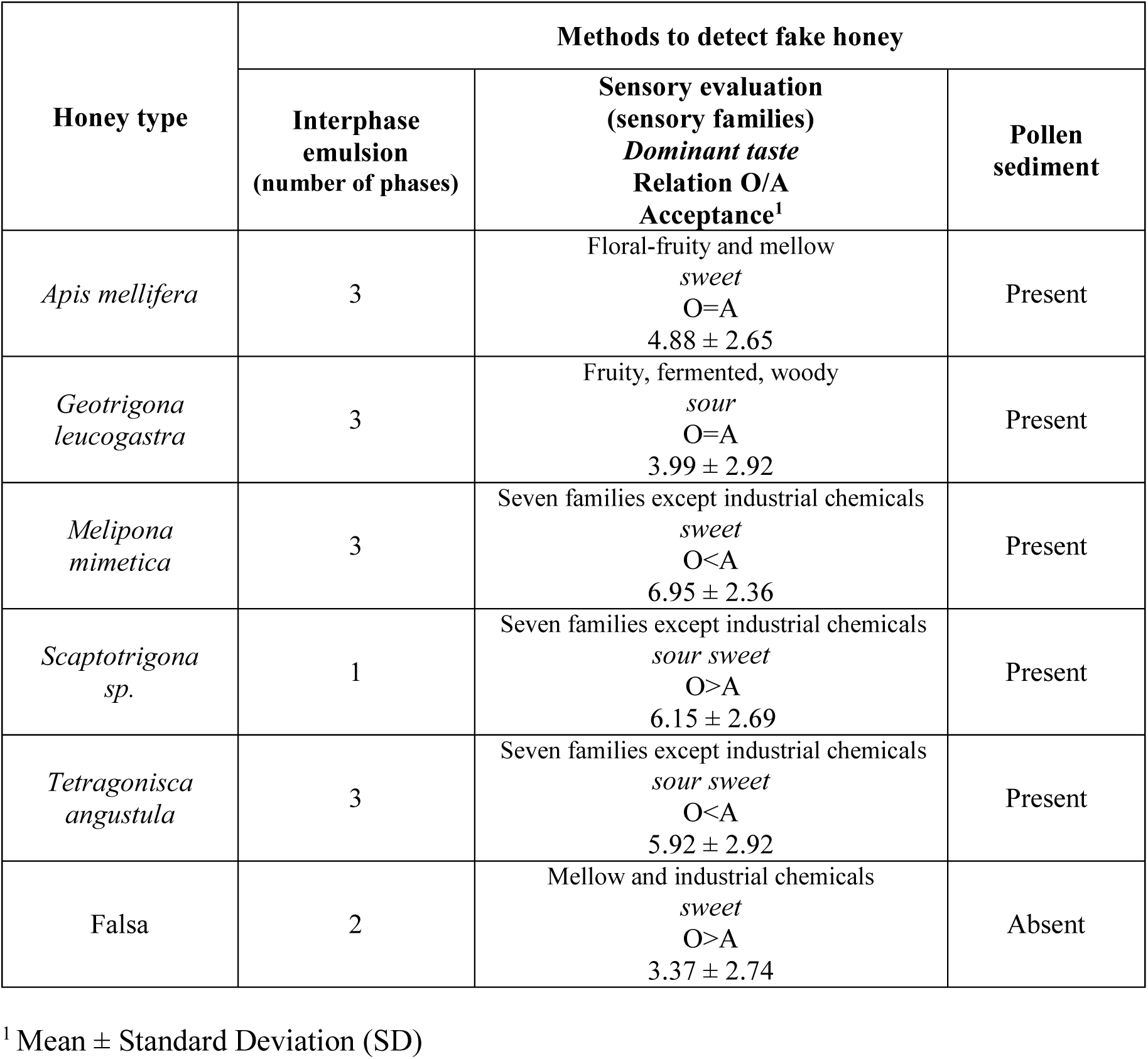
Three alternative methods to detect fake honey.

In the interphase emulsion test, the false honey presented two phases, unlike the genuine honeys of *Apis mellifera* and of stingless bees, with three phases, except *Scaptotrigona* one phase. The diversity of odor-aroma descriptors is greater in honey from stingless bees, *Apis mellifera* is represented by the floral-fruity and mellow families, and false honey by the mellow and industrial chemicals families. Acceptance varied between the lowest value of 3.37 for false honey < *Geotrigona* < *Apis* < *Tetragonisca* < *Scaptotrigona* and the highest value of 6.95 for *Melipona* honey. The dominant taste for each honey type varied among sour (*Geotrigona*), sweet (*Apis, Melipona* and fake) and sour-sweet (*Scaptotrigona* and *Tetragonisca*). *Geotrigona* honey was the most persistent whereas fake and *Apis mellifera* honeys were the less persistent, *Melipona, Scaptotrigona* and *Tetragonisca* with intermediate values of intensity 2. The relationship odor/aroma varied in the six honey types. The intensities of odor and aroma were equal for *A. mellifera* but *Geotrigona, Melipona* and *Tetragonisca* were more intense in the mouth, whereas the *Scaptotrigona* and the fake honeys where more intense in the nose. This false honey was particular for its notable presence of artificial eucalyptus essential oil, which was perceived more as an odor in the nose than as a retronasal aroma.

The sensory results obtained allow comparison of honey, pot-honey and false honey. Honey was perceived with odors-aromas from the floral-fruity, woody and mellow families. The honeys of *Geotrigona leucogastra, Melipona mimetica, Scaptotrigona* sp. and *Tetragonisca angustula* were more complex with components in all families except the industrial chemicals family. For fake honey, only two sensory families, mellow and industrial chemicals, were perceived.

The pollen sediment was not obtained in the fake honey but it was obtained in the five types of genuine honey. Melissopalynology is based on the analysis of the pollen sediment of honey, which is impossible in fake honey without sediment. Filtered honeys to delay cristallyzation are not suitable for melissopalynological analysis either.

Two organoleptic discoveries are reported here for the honey and nest of the Ecuadorian ‘Catiana’: 1. The biosurfactant activity in the authenticity test applied to honey from *Scaptotrigona* sp. by visual observation, and 2. The smell of blue Roquefort-type cheese from the nest of *Scaptotrigona* sp. by olfactory observation.

Figure 1 shows a diagram with the results obtained for the different types of honey with the honey authenticity test by interphase emulsion. Here the decision making is very simple, the honey with two phases is false, the honey with three phases is genuine as indicated by Vit (1998), and the new pattern of having a unique phase is also a genuine honey of *Scaptotrigona* sp. The origin of this new result for the genuine honey of *Scaptotrigona* sp. with a phase, was an unsolved puzzle for long time. Images of the test for each type of honey analyzed are also illustrated, where these difference between the two types of genuine honey, with a unique emulsion phase dispersed in the reaction volume (*Scaptotrigona*) or with an interphase emulsion, intermediate between the aqueous and the ethereal phases (*Apis, Geotrigona, Melipona*, and *Tetragonisca*) are easily observed. Fake honey presented only two phases, one aqueous and one ethereal, without emulsion.

**Figure 1.**
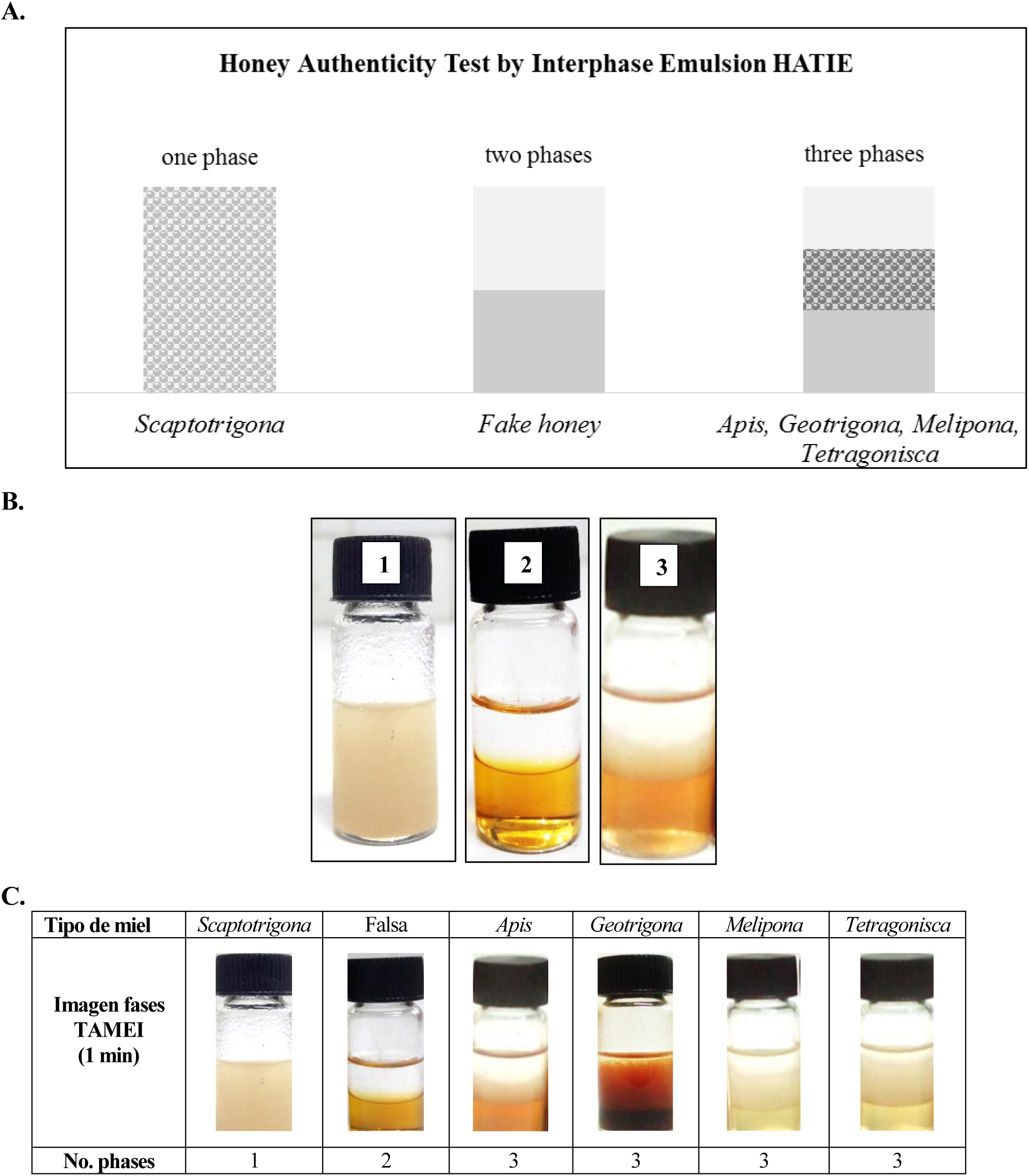
Number of phases formed with the HATIE **A**. Diagram with number of phases formed in the Honey Authenticity Test by Interphase Emulsion **B**. Genuine *Scaptotrigona* sp. honey has a biosurfactant that causes a single phase (first vial 1), there are two phases in the fake honey (second vial 2), and three phases in the genuine *Melipona* honey (third vial 3). **C**. Results in different honey types *Scaptotrigona* honey (one phase), 2. Fake honey (two phases), 3. *Apis, Geotrigona, Melipona, Tetragonisca* honeys (three phases)

A further HATIE screening was carried out with various Asian, Australian, European and Neotropical honey types (*Apis cerana, Apis dorsata, Apis mellifera*, unifloral, *Melipona fasciculata, Melipona scutellaris, Scaptotrigona depilis, Scaptotrigona mexicana, Tetragonisca angustula* and fake), which is presented in Table 2. The genuine honey presented one or three phases and the false honey two phases. *Scaptotrigona depilis* from Argentina, Bolivia, and Brazil, a meliponine honey from Malaysia informed to be *Geotrigona thoracica* without entomological authority, and that of *Tetragonula carbonaria* from Australia also presented one phase.

**Table 2.** HATIE screening of global honey types

| Bee<br>"Ethnic name" | Interphase<br>emulsion<br>(number of phases) | Botanical origin | Country |
| --- | --- | --- | --- |
| *Apis cerana* | 3 | - | Japan |
| *Apis dorsata* 'Tualang' | 3 | Tropical forest | Malaysia |
| *Apis mellifera* | 3 | Lavander *Lavandula officinalis* LAMIACEAE | France |
|  | 3 | Persian manna *Astragalus* FABACEAE | Iran |
|  | 3 | Chestnut *Castanea sativa* FAGACEAE | Italy |
|  | 3 | Orange *Citrus* RUTACEAE | Japan |
|  | 3 | Acacia *Acacia mangium* FABACEAE | Malaysia |
|  | 3 | 'belimbing' tamarindo chino *Averrhoa carambola* OXALIDACEAE | Malaysia |
|  | 3 | Durian *Durio ziberthinus* MALVACEAE | Malaysia |
|  | 3 | 'gelam' *Melaleuca cajaputi* MYRTACEAE | Malaysia |
|  | 3 | 'getah' Caucho *Hevea brasiliensis* EUPHORBIACEAE | Malaysia |
|  | 3 | 'kelapa' Coco *Cocus nucifera* ARECACEAE | Malaysia |
|  | 3 | 'nanas' Piña *Ananas comosus* BROMELIACEAE | Malaysia |
|  | 3 | Rambutan *Naphelium lappaceum* SAPINDACEAE | Malaysia |
|  | 3 | Manuka *Leptospermum scoparium* MYRTACEAE | New Zealand |
|  | 3 | Pohutukawa *Metrosideros excelsa* MYRTACEAE | New Zealand |
|  | 3 | Rewarena *Knightia excelsa* PROTEACEAE | New Zealand |
|  | 3 | - | Venezuela |
|  | 3 | Acacia *Robinia pseudoacacia* FABACEAE | Czech Republic |
|  | 3 | Strawberry *Rubus idaeus* ROSACEAE | Czech Republic |
|  | 3 | Sunflower *Helianthus annuus* ASTERACEAE | Czech Republic |
|  | 3 | Rape Brassica napus BRASSICACEAE | Czech Republic |
|  | 3 | Linden *Tilia cordata* MALVACEAE | Czech Republic |
| *Frieseomelitta paupera*<br>'Guanotica' | 3 | - | Venezuela |
| *Geniotrigona thoracica*<br>'Kelulut' | 1 | - | Malaysia |
| 3 *Melipona fasciculata*<br>'Tiúba' | 3 | - | Brazil |
| 8 *Melipona scutellaris*<br>'Uruçu' | 3 | - | Brazil |
| *Scaptotrigona depilis*<br>'Tapezuá' | 1 | - | Argentina |
| *Scaptotrigona depilis*<br>'Obobosí' | 1 | - | Bolivia |
| *Scaptotrigona depilis*<br>'Mandaguari' | 1 | - | Brazil |
| *Scaptotrigona mexicana*<br>'Pisil Nekmej' | 3 | - | Mexico |
| *Tetragonisca angustula*<br>'Jataí' | 3 | - | Brazil |
| *Tetragonisca angustula*<br>'Mariola' | 3 | - | Costa Rica |
| *Tetragonisca angustula*<br>'Ramichi' | 3 | - | Peru |
| *Tetragonisca angustula*<br>'Angelita' | 3 | - | Venezuela |
| *Tetragonula biroi*<br>'Kiwot' | 3 | - | Philippines |
| *Tetragonula carbonaria*<br>'Karbi' | 1 | - | Australia |
| Fake | 2 | - | Brazil |
|  | 2 | - | Colombia |
|  | 2 | - | Malaysia |
|  | 2 | - | Singapore |
|  | 2 | - | Venezuela |

Honeys of *Apis cerana* from Japan and *Apis dorsata* ‘Tualang’ –forest from Malaysia– *Apis mellifera* unifloral of acacia, *Acacia mangium* FABACEAE, ‘belimbing’ star fruit *Averrhoa carambola* OXALIDACEAE, durian *Durio ziberthinus* MALVACEAE, ‘nanas’ pineapple *Ananas comosus* BROMELIACEAE, ‘gelam’ *Melaleuca cajaputi* MYRTACEAE, ‘getah’ rubber tree *Hevea brasiliensis* EUPHORBIACEAE, ‘kelapa’ coconut *Cocus nucifera* ARECACEAE, rambutan *Naphelium lappaceum* SAPINDACEAE and ‘Kelulut’ –stingless bees– *Geniotrigona thoracica* of Malaysia were tasted during a Workshop on Sensory Evaluation of Asian Honeys 26 and 27 September before the *11th Asian Apicultural Association’s Conference & ApiEXPO*, Kuala Terengganu, Malasia, 28 Septiembre - 2 Octubre 2012. *Frieseomelitta paupera* ‘Guanotica’ from Venezuela, *Melipona fasciculata* ‘Tiúba’ y *Melipona scutellaris* ‘Uruçu’ from Brazil three phases. One phase was observed in the honey of *Scaptotrigona depilis* ‘Obobosí’ from Bolivia, ‘Tapezuá’ from Argentina and ‘Mandaguari’ from Brazil. Three phases were formed in the honey of *Scaptotrigona mexicana* ‘Pisil Nekmej’ from Mexico, *Tetragonisca angustula* ‘Jataí’ from Brazil, Mariola from ‘Costa Rica’, ‘Ramichi’ from Peru and ‘Angelita’ from Venezuela, and *Tetragonula biroi* ‘Kiwot’ from Philippines. *Tetragonula carbonaria* ‘Karbi’ from Australia formed one phase. Unifloral *Apis mellifera* honeys of lavender *Lavandula officinalis* from France, *Astragalus* from Iran, chestnut of *Castanea sativa* from Italy, manuka *Leptospermum scoparium*, pohutukawa *Metrosideros excelsa* and rewarena *Knightia excelsa* from New Zealand, acacia *Robinia pseudoacacia*, raspberry *Rubus idaeus*, sunflower *Helianthus annuus*, rape *Brassica napus*, and linden *Tilia cordata* from Czech Republic, developed three phases. Fake honeys from Brazil, Colombia, Malaysia, Singapore and Venezuela, formed two phases with the HATIE. All these botanical and entomological origins were informed, some were in the honey labels, but vouchers of botanical and entomological origins were not available.

## 4. Discussion

### 4.1 Honey Authenticity Test by Interphase Emulsion

Fake honeys were detected by this test because they do not form an intermediate emulsion as they do not have the characteristic components of genuine honey (Vit, 1998). This behavior was observed in 500 Venezuelan commercial honeys (Vit et al., 1994) where the idea of the interphase test arose, since genuine honeys generate three phases and false honeys only two. It is important to differentiate interface, which refers to the contact surface, from interphase, which refers to the volume of the emulsion (Geckeler et al., 1997). However, Table 1 shows that the honey from *Scaptotrigona* sp. produces a unique phase. This difference is illustrated in the diagram of Figure 1A. It is a cloudy phase as can be seen in Figure 1B. With the light microscope, this phase (vial 1) consists of multiple bubbles of liquid-air interaction at 400X, which contract and collapse on the slide, like the interphase (vial 3). The interphase emulsion that separates the ethereal phase from the aqueous phase is not formed, but dispersed in the reaction volume as a result of the action of a biosurfactant, therefore a single phase is observed. It is a phenomenon of the nest of this species of *Scaptotrigona* – not observed in honeys from *Apis mellifera, Frieseomelitta, Geotrigona, Melipona, Nannotrigona*, or *Tetragonisca* from Argentina, Bolivia, Brazil, Colombia, Costa Rica, Ecuador, Mexico, Peru, Trinidad & Tobago and Venezuela-

How to explain this differentiated interphase phenomenon? Biosurfactants produced by a yeast associated with this bee but not with the others could originate the phase observed only in *Scaptotrigona* sp. honey. There are interesting associations of microbes with insects (Schmidt and Engel, 2021). Bees function as vectors to transport yeasts from their natural reservoirs in floral nectar to the different substrates of bee nests (Starmer and Lachance, 2011). The yeast *Candida bombicola* initially isolated from *Bombus* bumblebee honey synthesizes a sophoroside with surfactant properties (Kim et al., 1997), in particular extracellular glycolipids with biosurfactant activity, known as sophorolipids C34H58O15. Sophorolipids (SL) are composed of a sophoroside molecular fraction (glucose disaccharide with an unusual β-1,2 bond whose etymology derives from *Sophora japonica* pods) and a lipid tail that is the aglycone.

The nomenclature of the yeast *Candida bombicola* changed with the new genus, and since 1998 it has been known as *Starmerella bombicola* (Rosa and Lachance, 1998); however, *Candida bombicola* still appears in some publications (C.A. Rosa, personal communication, 7^th^ February 2022). In other cases gender change is indicated as in *Starmerella* (*Candida*) *bombicola* JCM 9596 from the Japan Microorganism Collection at the Riken BioResource Center (Hirata et al., 2021). The study of the surfactants produced by *Starmerella* is of current interest due to its biotechnological applications (Van Renterghem et al., 2018). SLs are the surfactants of the future due to their versatility in formulations and because they are ecologically friendly. In fact, 75% of the research on *Starmerella bombicola* focuses on its SLs. But why does *S. bombicola* invest so much energy to produce this secondary metabolite? Although SLs do not represent osmotic protection in common habitats such as nectar and honey, they protect the niche of *S. bombicola* with a double advantage to compete against other microbes: 1. Extracellular storage of usable energy in starvation emergency, and 2. Antimicrobial activity (De Clercq et al., 2021). Proteomic and genetic research on *S. bombicola* is recent, such as the study of its exoproteome (Ciesielska et al., 2014) and its transportome (Claus et al., 2022), demonstrating the potential of omics sciences for the functional design of molecular biosynthesis in industrial biotechnology. One could think of some species of *Starmerella* or the same *S. bombicola* associated with *Scaptotrigona* sp. from Ecuador, which should be isolated from their honey to demonstrate this potential microbial origin of a biosurfactant that causes the behavior observed in the honey authenticity test based on an interphase emulsion.

*Starmerella bombicola* was identified in the honey microbiome of *Scaptotrigona bipunctata* and *Scaptotrigona depilis* from Brazil, but was absent in *Scaptotrigona tubiba* and other 14 species of stingless bees studied (Echeverrigaray et al., 2021). What are the traits of these two species of *Scaptotrigona* for the yeast *Starmerella bombicola* to be associated with them? They could be related to the bee diet, secretions, or even microbiome. The result of the interphase emulsion test performed with *Scaptotrigona depilis* honey from Argentina, Bolivia and Brazil, was one phase (See Table 2), which could confirm the production of extracellular sophorolipids with biosurfactant activity by a yeast type *Starmerella bombicola*, located in the same subclade. The honey produced by the Australian *Tetragonula carbonaria* and the Malaysian ‘Kelulut’ putatively *Geotrigona thoracica*, also had a one phase result. Biosurfactant sophorolipids are produced by multiple species of the yeast clade, subclade such as *Candida apicola, Candida riodocensis, Candida stellata* and a new species Candida sp. NRRL Y-27208 (Kurtzman et al., 2010).

The presence of sophorosides has also been associated with the botanical origin of bee and floral substrates, they are not SL but flavonoid sophorosides. For example: 1. Rosemary nectar extracted from the stomach of *Apis mellifera* contains glycosides, 93% kaempferol 3-sophoroside and 7% quercetin 3-sophoroside, but only their aglycones were found in honey (Gil et al., 1995), 2. Carnations contain kaempferol 3-O-sophoroside (Iwashina et al., 2010), 3. Bee pollen contains quercetin 3-O-sophoroside (Čeksteryte et al., 2016).

### 4.2 Sensory evaluation of honey

The participation of people as data generators of sensory analysis, makes this discipline influenced by factors such as social, cultural, religious, psychological, and climatic factors, physical status of the individual, availability, and nutritional education (Jain and Gupta, 2005). Sensory evaluation of honey derives from its chemical composition, its perception by sensory receptors, and cognitive interactions to discriminate (acceptance scale), quantify (intensity scale) and describe (vocabulary) its sensory attributes. Sugars confer sweetness, and aliphatic organic acids (AOA) in addition to acidity impart particular odors and aromas. Gluconic acid is dominant in honey from *Apis mellifera* (Stinson, 1960), but other AOA such as acetic acid and lactic acid predominated in honeys produced by *Geotrigona, Melipona*, and *Scaptotrigona* (Vit et al., submitted). These AOA are never isolated in nature, and their combinations are of industrial interest in order to enhance the flavor of foods (Dziezak, 2016). Knowing the sensory properties of AOA is very useful. In the first study conducted with gluconic acid and acetic acid monomers, a significant correlation of their proportions with the kombucha flavor score was demonstrated (Li et al., 2022).

In a previous study with an Ecuadorian Kichwa panel, the thick honeys (*A. mellifera* and fake) were clearly separated from the less dense stingless bee honeys (*Geotrigona, Melipona* and *Scaptotrigona*) by the Free Choice Profile (FCP). Each assessor perceived a different number of the 28 sensory attributes obtained for these honeys (Vit et al., 2017). The lexicon developed to describe honeys by descriptive quantitative analysis (Vit, 1993; Anupama et al., 2003; Persano Oddo and Piro, 2004; Galán-Soldevilla et al., 2005) was similar to the descriptors used in the (FCP) of meliponine honeys from Brazil (Ferreira et al., 2009).

The similarities and differences found between the sensory descriptors of the six honey types studied here –genuine honey from five entomological origins and fake honey– are valuable information for the normative scope of the Codex Alimentarius Commission. The proposal of a honey standard for each genus of stingless bees is justified, such as the first standard for *Melipona* honey, from the state of Bahia in Brazil (ADAB, 2014), and the proposal made for *Scaptotrigona* honey (Vit, 2017). In addition to routine physicochemical parameters in honey quality standards, sensory evaluation provides discriminating elements to differentiate genuine honey from fake honey, and honey produced by different species of bees. Additionally, sensory evaluation is a direct way to perceive metabolites of microbial origin in honey, without the need for large analytical equipment. This was demonstrated in the Sensory Evaluation Workshop of Stingless Bee Honey, Mérida, Venezuela (2007), with experts from Brazil, Colombia, Guatemala, Panama and Venezuela, where the Fermentation sensory family was proposed. The corresponding sensory descriptors for the proposed subfamilies: 1. Acetic, 2. Alcoholic, and Lactic (Vit, 2008a), are metabolites of the microbiota associated with stingless bees, with precise functions in the bee colony. Both the nutritional improvement of the substrate, the defense against entomopathogens, and others such as: 1. Acetic acid and lactic acid bacteria (Leonhardt and Kaltenpoth, 2014), 2. Yeasts (Rosa et al., 2003; Echeverrigaray et al., 2021), and 3. Edible filamentous fungi Monascus sic *Zygosaccharomyces* grown by *Scaptotrigona depilis* in brood cells (Menezes et al., 2015), orchestrated with two other nest microbes –*Candida* sp. and *Monascus ruber*– that modulate metamorphosis from larva to pupa (Paludo et al., 2019). Sensory evaluation brings us closer to the evolution of the associations between microbes and bees, to their ecological host-host role as investigated by Schmidt and Engel (2021), in mutualisms possibly originating from the Late Cretaceous, the era of the first fossil of a bee in our planet: *Cretotrigona prisca*, a stingless bee (Michener and Grimaldi, 1988a,b; Camargo, 2013). Some aromas of honey are so harmoniously amalgamated that if we knew how to express these sensory attributes with musical notes, it could be more precise.

In his taxonomic notes on *Scaptotrigona*, Engel (2022b) is surprised that in Rasmussen and Cameron’s (2010) phylogenetic study, *Scaptotrigona* was not placed close to the shy *Nannotrigona*, but in a clade with *Oxytrigona* Cockerell, with a higher degree of aggressiveness –which I know because when I collected my first Ecuadorian pot-honey I had to remove those bees from my hair with water– For this reason the scaptotrigonicultors work with a veil. Engel specifies that *Nannotrigona* and *Scaptotrigona* are morphologically similar, but have other non-morphological features that separate them. Just as *Scaptotrigona* is morphologically more similar to *Trigonini* than to *Melipona*, but in the chemical composition of its honeys, it turned out to be more similar to *Melipona* (Vit et al., 1998). Remarkable phylogenetic demonstrations could emerge by broadening the spectrum of biological traits not limited to morphology. Metabolites of microbial origin are novel for interpreting coevolution with stingless bees and the underlying causes of evolutionary preferences among bee-microbe associations. The study of the bacterial, fungal and yeast microbiomes associated with different species of stingless bees and their substrates begins to reveal particular differences between specialized and generalist dynamic evolution.

### 4.2 Microscopic observation of the pollen sediment

This test is very simple but requires a microscope, which is a basic equipment in bioanalysis but rare in food quality control laboratories. Fake honeys do not contain pollen, while genuine honeys contain pollen grains of different sizes, shapes, and exine surface structures. Yeasts, fungal mycelia and their spores are also visible. The difference between the presence and absence of pollen is enough to detect false honey.

### 4.3 Natural fermentation of pot-honey and studies of traditional fermented food

Pot-honey produced by stingless bees is a fermented product. Traditionally, fermented honey from *Apis mellifera* was considered a defect caused by harvests of unripe honey with high moisture content. Fake honeys meet the <20% moisture requirement. However, the honey produced by Meliponini can ferment from the nests of stingless bees, a fermentation that can continue after harvest (Drummond, 2013), and therefore merited the inclusion of the new sensory family Fermentation, with the acetic, alcoholic and lactic subfamilies, and their sensory descriptors: vinegar, pot pollen, brandy, fermented fruit, brewer’s yeast, liquor, must, sake, vinasse, white wine, red wine, miso, cheese, yogurt (Vit, 2008c). Fermentation is not a defect of meliponine honey but its method of food preservation. That is why there are so many yeasts in their nests (Rosa et al., 2003; Silva et al., 2019; Echeverrigaray et al., 2021; Gavazzoni et al., 2022) as in industrial fermentations. Ethanol production in *Tetragonisca angustula* honey from Venezuela was stabilized in 30 days post-harvest (Pérez-Pérez et al., 2007) and lactic acid bacteria were inactivated in *Heterotrigona itama* honey from Malaysia (Yaacob et al., 2021). In another study, the proportion of gluconic acid was optimized to improve the taste quality of kombucha, using a Synthetic Microbial Community (SMC). *Starmerella davenportii* with the highest ethanol yield and *Gluconacetobacter intermedius* with the highest acetic acid yield were selected to rebuild the microbial community that allowed obtaining a better sensory quality kombucha with 74% gluconic acid (Li et al., 2022).

### 4.4 The Roquefort blue cheese smell in the nest of Scaptotrigona sp

Lipolysis gives rise to carboxylic acids and other malodorous molecules that impart sensory complexity to fungus-inoculated blue cheeses, such as British Stilton, Italian Gorgonzola, or French Roquefort. For example, fatty acids form methyl ketones (alkan-2-one, heptan-2-one, nonan-2-one) with blue cheese notes, and they also react with alcohols such as ethanol to produce flavoring esters. Other decarboxylation, deamination, oxidation, and reduction transformations produce short-chain volatile organic compounds. Lactate and citrate produce molecules such as butter-flavored diacetyl, ethanal, and ethanol (Cotton, 2010). Explaining the origin of the sensory complexity of blue cheese in a dairy-free stingless bee nest is astounding, and a challenge soon to be revealed with the advancement of omic sciences in stark contrast to the growing cross-border bureaucracy, which far from cooperating, makes it difficult to transfer honey from the tropics to specialized laboratories. It can be said that the biotechnology of inoculum-type fungi in blue cheese produces this biotransformation in some nest substrate of *Scaptotrigona* sp. perceived by the sense of smell, although its functions for the bee colony are not known. It could also be an unknown secretion from the bee or a derivative from foraged materials, particularly biotransformed.

In a European patent, three aliphatic organic acids that cause unpleasant odors (acetic acid, butyric acid and isovaleric acid) are indicated as by-products of the microbial biosynthesis of sophorolipids (European Patent EP 2 821 495 B, 2016).

### 4.5 On fake honey, omic sciences and honey consumer protection

Fake honey could be recognized by sensory evaluation. Overheated honey for its production could exceed the maximum permitted amount of hydroxymethylfurfural (40 mg/kg) in the official standards, but if it contains corn syrup with high fructose content, HFCS type (high fructose corn syrup), it does not increase the content of hydroxymethylfurfural (HMF), and fake honeys would meet this requirement for the standard.

Walker et al. (2022) reviewed multiple analytical techniques needed to detect and report sophisticated honey adulteration. A recent strategy to detect fake honey is to apply metabolomics in order to identify foreign compounds in bee honey. Honey diluted in deuterated water allowed detection of benzoic acid, citric acid, sorbic acid and vanillin in high concentrations by ^1^H-NMR, as well as HMF and sucrose (Schievano et al., 2015). In Malaysia, 80% of the honeys are counterfeit adulterated and their packaging is very luxurious (personal observation). ^1^H-NMR spectra of genuine stingless bee honeys with controlled adulterations of 1% C3 and C4 sugars were fingerprinted for their detection, along with routine OPLS-DA chemometric models in metabolomics (Yong et al., 2022).

There are few molecular studies on the maturation mechanism of honey bees. Therefore, the metabolomics of the formation of mature honey is a resource to recognize false honeys (Guo, 2021). A state-of-the-art study compared immature honey with mature honey obtained during a turnip Brassica napus bloom by UPLC-QToF-MS-based metabolomics coupled with multivariate analysis of PCA, OPLS-DA, and VIP, and its authors found that mature honey operculada doubles the content of decenedioic acid originating from the bee (Sun et al., 2021).

Depending on the objective of the study of fake honey in the market, metabolomic approaches can be used to identify and quantify foreign substances to this product, either by NMR spectroscopy, by chromatographic or electrophoretic techniques coupled to multivariate statistics. These techniques are very effective for searching for markers of botanical (Schievano et al., 2016; Seraglio et al., 2021), geographic (Spiteri et al., 2015) and entomological (Vit et al., 2015a) origin.

It is recommended to identify the valuable bee products with Protected Geographical Indications (PGI) to value the agricultural product linked to its place of origin and also raise its quality and the reputation of honey professionals. These actions protect the consumer of honey by offering a genuine product. The Protected Designation of Origin (PDO) could be requested for honey from four types of stingless bees (*Geotrigona, Melipona, Scaptotrigona, Tetragonisca*). The IGP could be proposed for particular habitats, at least in its geographic stripes, coast, mountains, and jungle, in the case of Ecuador, and corresponding habitats in other countries. The Guaranteed Traditional Specialty (ETG) protects the production method, which is artisanal in the case of honey from Meliponinos “The PDO/IGP protects a name that identifies a product originating from a certain place, and the ETG protects the production methods and traditional recipes. In a product with PDO/PGI, the specificity is due to the origin of the product, while in one with ETG it is due to its traditional nature” (Mapa, mapa.gob.es).

### 4.6 A story about stingless bees, associated yeasts, and etymologies

I accepted a tribute from C.A. Rosa from the Universidade Federal de Minas Gerais in Brazil, to name a new species of yeast for my contributions in bee systematics and ecology – he knows I am a food scientist– An attentive and formal Email was received, with a manuscript that I could start to understand several years later. *Starmerella vitae* sp. nov. Santos, Lachance & C.A. Rosa, 2018, yeast isolated from the stingless bee *Trigona fulviventris* Guérin, 1844 known as ‘Jicote’, from Santa Rosa National Park, Guanacaste, Costa Rica, and from Brazilian flowers of the vine *Thunbergia grandiflora* ACANTHACEAE, from Ilha Grande State Park (Atlantic Tropical Forest), Rio de Janeiro state, and *Tabebuia* spp. BIGNONIACEAE Belo Horizonte, Minas Gerais state. D.W. Roubik of the Smithsonian Tropical Research Institute in Panama, and C.A. Rosa witnessed this food honey is transforming into ecology of microbiota and its metabolites in nests of meliponines. And Rosa wanted her to be accompanied by two special friends, the memory of João M.F. Camargo of the Universidade de São Paulo, who identified the meliponines from Venezuelan honeys, dedicated *Starmerella camargoi* sp. nov. Santos, Morais, Lachance & C.A. Rosa, 2020, and to the co-editor of meliponine cerumen pot-reactors to whom he dedicated *Starmerella roubikii* sp. nov. Lachance & C.A. Rosa, 2018. M.S. Engel of the Department of Ecology & Evolutionary Biology, Entomology Division, Natural History Museum, University of Kansas, Lawrence, is very active updating Meliponini taxonomies. With M.S. Engel, this honey from fermented foods ventures into the systematics of bees with an associated yeast for its elaboration, thus C.A. Rosa was a visionary and without realizing it, her baptism worked. In the phylogram based on the D1/D2 domains of the LSU ribosomal large subunit rRNA gene, *Starmerella vitae* is in the subclade of *Starmerella bombicola* with the bio-sophorolipid-producing species.

## 5. Conclusions and Recommendations

Three alternative methods were presented to verify if a bee honey is genuine or false, which were used in six Ecuadorian honeys. In order to detect false honey and protect the consumer. The interphase emulsion method formed by stirring ethyl ether with aqueous honey dilution is the fastest. The classic analysis such as those of the NTE INEN (2016) are between this alternative approach and that of the major analytical equipments (CE, HPLC, NMR). However, they are limited, and do not allow the identification of any foreign additive in the honey, like other national standards they do not detect all false honeys, and group genuine fermented or heated honeys as out of standard.

Melissopalynology starts with the observation of the pollen sediment. The presence of pollen grains in the sediment was an indicator of honey authenticity, although its botanical origin was not identified and the types of pollen were not quantified, it was evaluated that the sediment did not contain foreign particles, which were not detected in the genuine honeys analyzed. The absence of sediment observed in the fake honey was an indicator of honey manufactured with syrups or industrial sugars, having no pollen traces derived from nectar or honeydew raw materials used by honey making bees.

Sensory evaluation is a method used to value honey by its organoleptic attributes (smell, aroma, taste, persistence). In this research the objective was to differentiate genuine from fake honey, which was achieved. The wealth of types of tropical honey deserves to include its sensory evaluation together with its valorization of botanical, entomological and geographical origin. At the Stingless Bee Honey Sensory Evaluation Workshop, Mérida, Venezuela, 2007, the guiding thread was a poem by Alfa Bet that concluded as follows: …/ *art of kissing honey* / *melisophy walk* / *calmly* / *honey will kiss you*. And that I recommend. Continue to incorporate new sensory descriptors as they appear, are perceived and needed for sensory descriptions of honey processed by stingless bees in the cerumen pots of their nests where fermentations with associated microbiota occur. I hope that soon, in addition to the Fermentation family with metabolites of microbial origin, the Microbiota family can be proposed, with some O-A characteristics.

For the first time, the results of a unique phase in the Honey Authenticity Test of *Scaptotrigona* honey, was associated with the presence of a natural biosurfactant, possibly of microbial origin. The production of sophorolipids by the yeast *Starmerella bombicola* – frequent in the scientific literature– and its associations with Neotropical stingless bees, also widely demonstrated, are the basis of this interpretation. The microbiome of *Scaptotrigona* sp. from Ecuador has not been studied in the bee or in the different substrates of its nest, where the suspected microbial origin of the biosurfactant in this investigation could be evidenced. It will be necessary to study the nature and chemical functionality of this metabolite in another investigation, to postulate its functions for *Scaptotrigona* sp. and for the yeast that produces it. Likewise, the distinctive Roquefort cheese odor in its nest may have a microbial origin. I wanted to publish this contribution in Spanish because it is the honey of a Latin American bee, but it did not fit for Revista de Biología Tropical as a tribute to Don Alvaro Wille, Interciencia and Agroalimentaria. Their interactions are visible in the edited text expanded for biotechnology and social interest. The new interdisciplinary rurality of Pérez (2004) is ready to promote social development from territorial revaluation needed for all stingless bees kept in Ecuador. The ‘Catiana’ is embedded in the Ecuadorian geography and in the knowledge of those who care for their nests and harvest their honey. They are stingless bee keepers that would deserve a complex name –scaptotrigonicultors– imitating that of meliponicultors.

In addition, another application for this test was proposed, useful for those interested in honey metabolites. When a unique phase is obtained with this procedure, this test detects the presence of biosurfactant activity and becomes the Honey Biosurfactant Test (HBT). This use allows screening to select stingless bees associated with biosurfactant-producing microbiota in honey. In particular, if demonstrated, it could be a Test for Sophorolipids in Honey (SST) because the yeast *Starmerella bombicola* does not produce any biosurfactant but sophorolipids. Its industrial applications in food, bioremediation, cosmetics, pharmacy and ecological products for the home are known. The biotechnological development of biosurfactants of microbial origin has constant parametric optimizations of cell cultures (solid phase, use of dual lipophilic substrates, standardization). *Starmerella bombicola* is widely studied in biotechnology for its massive production of extracellular sophorolipids. It was initially isolated from the honey of the *Bombus* bumble bee, later from the stingless bee ‘Jicote’ *Trigona fulviventris* from Costa Rica. More recently from the honey produced by two species of stingless bees of the genus *Scaptotrigona* associated with this yeast in Brazil.

The type of emulsions obtained with the test deserves further research, and also the component causing the biosurfactant activity. The physical state of the *Scaptotrigona* honey is liquid. Is there any nanoemulsion given the biosurfactant activity observed under the test conditions? Just to think on possible applications of this honey itself. Nanoemulsions are nano-sized emulsions with medicinal industrial uses, manufactured for improving the delivery of active pharmaceutical ingredients (Jaiswal et al., 2015).

Knowing the types of genuine honey, characterizing them, and promoting their consumption with cutting-edge bee keeping and meliponiculture, targeting the informed consumer, is a shared commitment. The Ecuadorian National Institute for Standardization (INEN), the Phytosanitary and Zoosanitary Regulation and Control Agency (AGROCALIDAD) and the Ministry of Agriculture and Livestock (MAGAP) could join efforts to revitalize the future development of meliponiculture in Ecuador in all its provinces except Galapagos where the unique bee is the solitary *Xylocopa darwini* Cockerell, 1926.

Paradoxically, in this communication without microbiological analysis, the fungal presence was perceived from the nest of *Scaptotrigona* sp. by sensory observations different from the classic sensory evaluation of honey, and corresponding scientific interpretations: 1. Biosurfactant activity in honey perceived visually by a unique emulsion phase formed in the honey authenticity test by interphase emulsion for genuine honeys, and 2. Typical odor of blue cheese such as Roquefort obtained after inoculated fungi develop and perceived olfactorily when opening the nest.

## Ethics statement

The author declares that she has made the contributions that justify her authorship. There is no conflict of interest of any kind and the relevant ethical and legal requirements and procedures have been complied with no funding received.

## Acknowledgements

To the stingless bee keepers and bee keepers of the world, producers and defenders of genuine honey. To Carlos Augusto Rosa, Department of Microbiology, ICB, Universidade Federal de Minas Gerais, Belo Horizonte, MG, Brazil, for his long-standing research on microbial associations with stingless bees, and the many details harvested and shared on their metabolites. To my family for their constant and unconditional support, my parents Giovanna and Giovanni, my brothers Daniele, Massimo and Leo, all participated in the development of the honey authenticity test. The manufacture of the prototype was carried out in VITCA, Vanguardia de Industria Tecnológica C.A, Judibana, Falcón state, Venezuela. To Michael S Engel from Kansas University to see a new species in the specimen sent to Charles D Michener, and deposited at the Snow Entomological Museum Collection (SEMC –besides nearly 5 million pinned insect specimens–) where other collections saw dust or could not decipher. To Silvia R.M. Pedro, J.M.F. Camargo, and Carlos Vergara for their friendship, meliponine taxonomic talent, and timely availability. To the Scientific, Humanistic, Technological and Art Development Council of the Universidad de Los Andes, Mérida, Venezuela, for the trust and encouragement received since 1986 to investigate the honey produced by bees and their imitations. To the Universidad Técnica de Machala, El Oro, Ecuador, and the Prometeo Program. To the Universitá Politecnica delle Marche, Ancona, Italy.

